# Species and strain cultivation of skin, oral, and gut microbiota

**DOI:** 10.1101/2021.04.01.435439

**Authors:** Elizabeth Fleming, Victor Pabst, Amelia Hoyt, Wei Zhou, Rachel Hardy, Anna Peterson, Ryan Beach, Yvette Ondouah-Nzutchi, Jinhong Dong, Julia Oh

**Author notes:** These authors contributed equally to this work (Elizabeth Fleming, Victor Pabst). Corresponding author and lead contact: Julia Oh, Ph.D. The Jackson Laboratory for Genomic Medicine, 10 Discovery Drive, Farmington, CT.

## Abstract

Genomics-driven discovery of microbial species have provided extraordinary insights into the biodiversity of human microbiota. High resolution genomics to investigate species- and strain-level diversity and mechanistic studies, however, rely on the availability of individual microbes from a complex microbial consortia. Here, we describe and validate a streamlined workflow for cultivating microbes from the skin, oral, and gut microbiota, informed by metagenomic sequencing, mass spectrometry, and strain profiling.

## INTRODUCTION

Genomics-driven innovations, such as 16S ribosomal RNA (rRNA) sequencing and shotgun metagenomics have been powerful drivers of discovery in a wide range of microbial ecosystems, particularly humans. The blueprints created by large-scale studies have elucidated an extraordinary microbial biodiversity across individuals, geographies, ethnicities, disease states, and lifestyles. Such blueprints have been critical for baseline characterizations of different ecosystems and for generating hypotheses by correlative analyses with phenotypes of interest. The natural succession of understanding host microbiome interactions are mechanistic investigations, potentiated by the findings from metagenomic surveys, and actuated by the availability of substrates of interest – namely, the microbes themselves.

Obtaining microbial isolates from a sample of interest has multiple values. First, sequencing isolates provides the highest quality reference genome sequences; genome reconstructions from metagenomic data can result in incomplete and fragmented genomes, as closely related genomes, low abundance genomes, and highly complex communities pose significant computational challenges that hinder accurate reconstruction of function and biodiversity. Second, an isolate in hand allows experimentation and manipulation to understand its genetics, molecular and physiological mechanisms, and inter- and intraspecies interactions. Third, isolates allow precise investigation of subspecies, or strain diversity, which is critical as individual strains of a microbial species can exhibit widely diverse phenotypes. For example, most *Escherichia (E.) coli* strains are commensals in the human gastrointestinal tract, while some strains can cause severe disease^1^. In addition, isolates are necessary for inferring evolutionary dynamics, transmission, and lineage tracking during infectious disease outbreaks^2^. Finally, a tremendous genetic and phenotypic diversity is encoded at the strain, rather than higher taxonomic levels, and individual disease susceptibility or severity phenotypes can be attributed to shared species, but unique strains. Different algorithms have been developed to infer strain diversity from metagenomic datasetse.g.,^3-5^, but these can vastly underestimate strain diversity. Correspondingly, approaches to recover isolates from microbiota would benefit significantly from accounting for strain diversity.

Finally, systematic methods for cultivation and recovery of microbial isolates from a sample is complicated by the notable body-site specificity in the human microbiome. For example, the gut harbors the highest biodiversity, with characteristic bacteria from Bacteroides and varied lactobacilli, enterobacilli and enterococci, *Bifidobacteria, Clostridia*, and methanogens. The oral cavity is typically populated with streptococci, *Haemophilus, Prevotella, Veillonella* genera, and the skin staphylococci, *Corynebacterium* and *Cutibacterium*^6^. Even within each of these body sites, significant local variation exists, such as the stomach vs. the small intestine vs. the cecum, oral pockets vs. the tongue dorsum, or the moist, oily, dry, or foot sites of the skin.

Numerous approaches have been defined to systematically cultivate microbes from different ecosystems, with a focus particularly in the gut. Extraordinary efforts have been made to increase the recovery of gut microbial biodiversity from this anaerobic environment, using up to 212 different culture conditions^7,8^, which might include a wide variety of different nutritive conditions or additives, different gas fractions, temperatures, pH, or inhibition via antimicrobials. Microfluidics devices^9^ to optimize and isolate cells, or metagenomic prediction of membrane epitopes for synthetic design of antibodies have also been used to capture microorganisms of interest^10^. Multiple, sequenced large-scale gut microbial culture collections have been recently established^8,11-13^, and these efforts have correspondingly increased the accurate annotation of metagenomic datasets. In contrast, while skin cultivation methods were prolific in the 1950s^14^, there was no potential to inform recovery using metagenomic characterizations, and fewer consolidated and systematic efforts exist for human oral or skin microbial cultivation, with recent efforts primarily targeted efforts to recover microbes of interest, like *Cutibacterium (C.) acnes*^15^ or Gram negatives^16^.

Here, our ultimate goals were to define a set of user-friendly cultivation conditions that would allow us to cultivate dominant microbiota from many different individuals and body sites, estimate recovery based on metagenomic data, and importantly, identify rapid, low-cost approaches to delineate strain diversity to better inform isolate selection for more laborious and costly whole genome sequencing. We cultivated isolates from the human gut, oral cavity, and two physiologically diverse skin sites on a streamlined set of different nutritive conditions and performed shotgun sequencing of matched samples to evaluate recovery. Finally, we report on the utility of Fourier Transform Infrared (IR) Spectroscopy to rapidly classify and type strains of common species. Taken together, this work builds on and consolidates approaches for generating culture collections from a variety of different environments, enabling a range of follow-up genomic and phenotypic characterizations.

## MATERIALS AND METHODS

### Sample acquisition

#### Ethics approval and consent to participate

Forehead and toeweb swabs (skin) and inner cheek and tongue dorsum (oral) swabs, and stool samples were obtained from our biorepository (IRB approved by the Jackson Laboratory Institutional Review Board), representing 25 different samples obtained from 12 individuals. For skin and oral microbiota, sites were swabbed rigorously using 2 PurFlock Ultra buccal swabs (Puritan Medical Products) for each site for thirty seconds before one swab was submerged into a 1.5mL Eppendorf tube containing 500μl R2A broth (R2A) culture media (Lab M) and the other one into a microfuge tube containing 350 μL Tissue and Cell Lysis buffer (Epicentre) and 100 μg 0.1 mm zirconia beads (BioSpec Products) for mWGS sequencing. Stool was self-collected using from volunteers using a BioCollector fecal collection kit (The Biocollective, Denver, CO). Volunteers collected the sample at home, adding a portion of the sample to an Omnigene gut tube (DNA Genotek) for preservation for sequencing prior to sending the sample. Upon receipt, samples were immediately frozen at −80 for storage. For WGS sequencing, approximately 50 microliters of the thawed Omnigene preserved stool sample was added to a microfuge tube containing 350 μL Tissue and Cell lysis buffer (Epicentre) and 100 μg 0.1 mm zirconia beads (BioSpec Products).

### Cultivation conditions (Table 1)

#### Skin and Oral

The 1.5mL Eppendorf tube containing the culture media was shaken thoroughly with the swab using a vortex mixer, and then diluted to the 1:100 and 1:1000 dilution to increase the chance of recovering single colonies. 50 μL from each dilution was then plated on half of an agar plate for each of our cultivation conditions.

#### Stool

The stool sample was thawed and approximately 5g was added to a 50 ml conical tube containing 15 ml PBS (for aerobic culture) and 15 ml deoxygenated PBS with 0.1% cysteine (for anaerobic culture) vortexed well for 5 minutes and left to settle for 15 minutes.

#### Direct plating

For the anaerobic samples, serial PBS/cysteine dilutions of 1:10,000 and 1:100,000 were plated on GMM plates and left to incubate at 37°C degrees in the anaerobic chamber for 48-72 hours until colony formation was observed.

#### Blood culture-assisted cultivation

To select for the growth of underrepresented and slow-growing species, the PBS/stool mixture was added to a variety of culture conditions and incubated for 3, 7, and 14 days prior to being diluted in PBS and plated on blood agar plates (**Table 1** and **Figure 1**). Plates were incubated in the atmosphere and temperature of the original culture for 24-72 hours until colony formation was observed.

**Table 1.** Cultivation conditions

**Skin cultivation conditions**
| **Media** | **Conditions** | **Origin** |
| --- | --- | --- |
| LB agar | Aerobic, Anaerobic, 37C | Thermo Fisher Scientific |
| R2A agar | Aerobic, Anaerobic, 37C | Thermo Fisher Scientific |
| TSA with 5% sheep blood | Aerobic, Anaerobic, 37C | BD Biosciences |
| Brucella agar | Aerobic, Anaerobic, 37C | AnaerobeSystems |
| BCYE agar | Aerobic, Anaerobic, 37C | Hardy Diagnostics |
| MacConkey agar | Aerobic, Anaerobic, 37C | Hardy Diagnostics |

**Oral cultivation conditions**
| **Media** | **Conditions** | **Origin** |
| --- | --- | --- |
| LB agar | Aerobic, Anaerobic, 37C | Thermo Fisher Scientific |
| R2A agar | Aerobic, Anaerobic, 37C | Thermo Fisher Scientific |
| TSA with 5% sheep blood | Aerobic, Anaerobic, 37C | BD Biosciences |
| Chocolate agar | Aerobic, Anaerobic, 37C | Hardy Diagnostics |
| Selective Strep agar | Aerobic, Anaerobic, 37C | Hardy Diagnostics |

**Stool cultivation conditions**
| **Culture device prior to plating onto TSA with 5% sheep blood** | **Conditions** | **Origin** |
| --- | --- | --- |
| Direct plating onto GMM agar | Aerobic-28°C |  |
| TSB (3, 7, 14d) + sheep blood (9% final vol) | Aerobic-37°C | BD Biosciences |
| Aerobic blood bottle (3, 7, 14d) + rumen fluid (9% final vol) | Aerobic-37°C | BD Biosciences |
| Aerobic blood bottle (3, 7, 14d) + sheep blood (9% final vol) | Aerobic-37°C | BD Biosciences |
| Aerobic blood bottle (3, 7, 14d); sample filtered at 5µm | Aerobic-37°C |  |
| BHI (3, 7, 14d) + vancomycin + colistin (10µg/mL each) | Aerobic-37°C |  |
| TSB (3, 7, 14d) | Anaerobic- 37°C |  |
| Anaerobic Blood bottle (3, 7, 14d) + sheep blood (9% final vol) | Anaerobic- 37°C | BD Biosciences |
| Anaerobic Blood bottle (3, 7, 14d) + rumen fluid (9% final vol) | Anaerobic-37°C | BD Biosciences |
| Anaerobic Blood bottle (3, 7, 14d); sample filtered at 5µm | Anaerobic-37°C | BD Biosciences |
| Anaerobic Blood bottle (3, 7, 14d); thermic shock (85C, 20 min) | Anaerobic-37°C | BD Biosciences |
| BHI (3, 7, 14d) + vancomycin + colistin (10µg/mL each) | Anaerobic-37°C |  |
| TSB (3, 7, 14d) + sheep blood (9% final vol) | Anaerobic-37°C |  |

**Figure 1.**
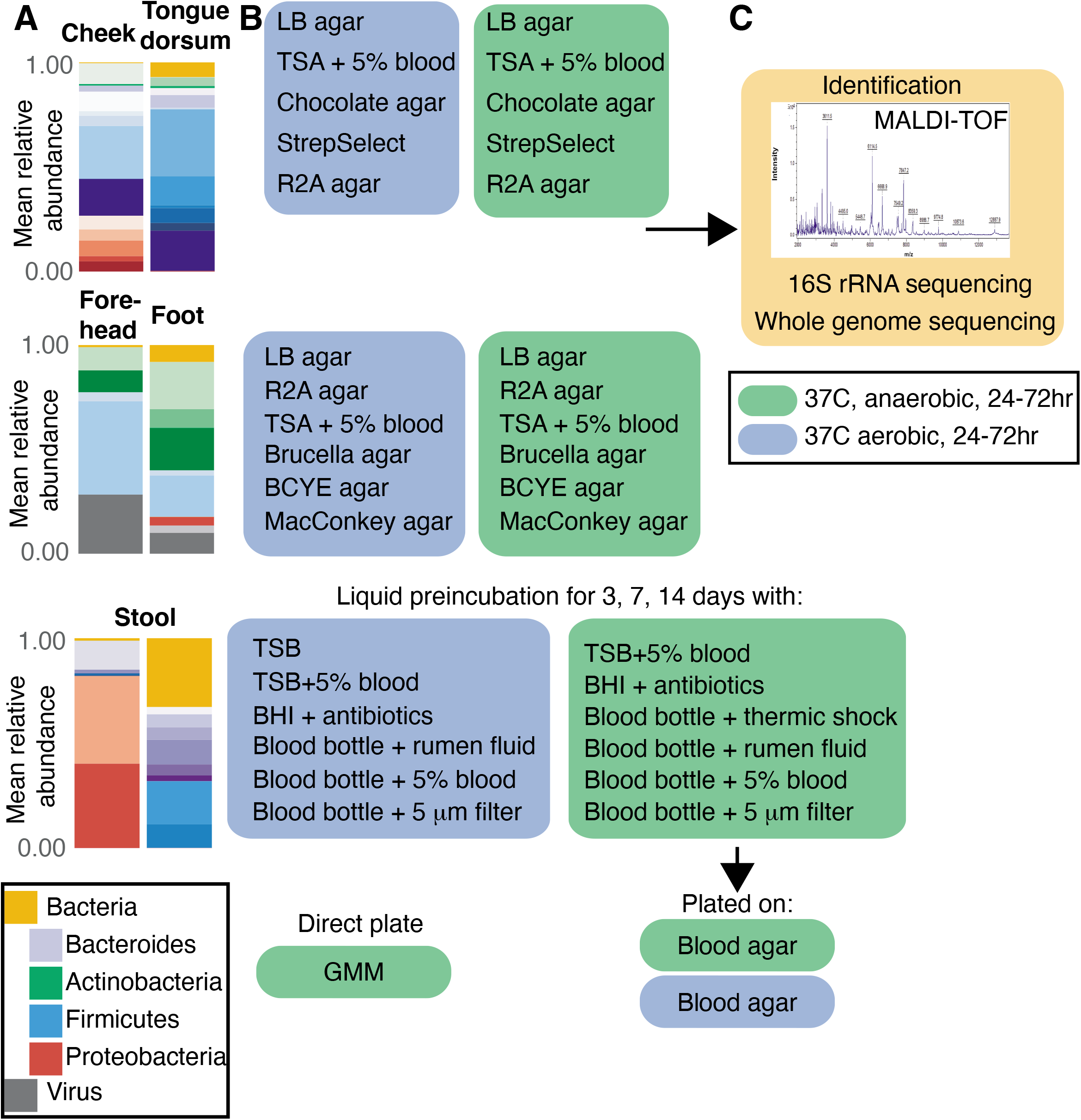
Culturomics pipeline. **A)** Metagenomic data were generated for each oral, skin or stool sample. Example relative abundance plots of major species are shown with colors corresponding to phylum as in the legend. Samples were then **B)** diluted and cultivated in a defined set of anaerobic (green boxes) and aerobic (blue boxes) cultivation conditions for oral, skin, and stool. After a defined period of incubation, individual colonies were picked, subcultured for purity, then **C)** identified using MALDI-TOF, 16S rRNA sequencing, or whole genome sequencing.

### Isolate identification

#### MALDI-TOF

We used matrix assisted laser desorption ionization-time of flight (MALDI-TOF, Bruker Daltonics, Germany) mass spectrometry to identify isolates. Ten to twelve single colonies from each cultivation condition were picked and replated onto a new blood agar plate, then grown for 24-48 hours to generate sufficient material for identification and archiving. Using a sterile transfer device (Puritan), bacteria were directly transferred to a MALDI target spot. We then used the ‘extended direct method’ for sample preparation, in which 70% formic acid is used to solubilize the bacterial cell wall prior to addition of matrix. One spot on the target was reserved for the bacterial test standard for calibration. Mass spectrometry analysis was performed using Flex Control 3.4 software (Bruker Daltonics, Germany). Colonies not recognized by MALDI-TOF were processed using the ‘protein extraction method, or by 16S rRNA sequencing, below.

#### Protein extraction

Bacteria failing identification by the extended direct method were first extracted. 10 mg of biological material (generous scoop of a 1 μL inoculation loop) was thoroughly suspended in 300 μL of High-Performance Liquid Chromatography water (Sigma-Aldrich) then thoroughly mixed with 900 μL 100% ethanol in a 1.5 mL Eppendorf tube. The tube was then centrifuged at 13,000 rpm for two minutes and the supernatant decanted. The tube was centrifuged again, and remaining ethanol was removed with a small pipet or allowed to air dry at room temperature. 10 μL 70% formic acid was thoroughly mixed to the pellet by pipetting, followed by 10 μL acetonitrile, then centrifuged for another two minutes. 1 μL of supernatant was used on each MALDI target spot, dried, and overlayed with 1 μL matrix. Mass spectrometry analysis was performed as above.

#### 16S rRNA sequencing

Bacteria failing identification by mass spectrometry and protein extraction were identified by 16S rRNA gene sequencing. Alkaline lysis was used to generate microbial DNA for PCR using universal primers 8F and 1391R (Turner, 1999). Sanger sequencing (GeneWiz, Inc) results were analyzed using BLAST (blast.ncbi.nlm.nih.gov). An identity score of 99% or higher was the threshold used for accurate species identification.

For archiving, the recovered bacteria were grown in tryptic soy broth growth media supplemented with 0.1mg/L Vitamin K and 5mg/L hemin 24-48 hours in 96 well plates at 37°C degrees in the appropriate atmosphere, then stored in 20% glycerol at −80°C degrees.

#### Whole genome sequencing

Rapid DNA extraction from *S. epidermidis* isolates was adapted from Köser et al. (2014). Briefly, 1 mL of overnight culture was centrifuged at 20,000 x g for 1 minute before the bacterial pellet was resuspended with 100 μL of 1X TE and transferred to a 2 mL bead beating tube with 100-125 μL 0.5 mm diameter glass beads (BioSpec Products). An additional 100 μL of 1X TE was added to the tube, followed by vortexing of the sample for 30 seconds at max speed (3000 rpm) on a vortex adapter (Mo Bio Laboratories). The mixture was then centrifuged at 13,000 x g for 5 minutes to pellet the cellular debris, and the supernatant was transferred to a new tube to be used as template for Nextera XT library preparation.

Sequencing libraries were made according to the Illumina standardized protocol using the Nextera XT DNA sample preparation kit (Illumina Inc.). All DNA samples were quantitated by Qubit HS (ThermoFisher Scientific) and diluted to 160pg/μl. The dual indexed paired-end libraries of genomic DNA were made with an average insert size of 400bp by taking 200pg DNA of each sample in optimized quarter reaction protocol, where all reagents for library preparation were taken in 1/4th amount. Tagmentation and PCR reactions were carried out according to the manufacturer’s instructions. The resulting Nextera WGS libraries were then sequenced with 2X150bp paired end reads on an Illumina HiSeq2500.

Sequencing adapters and low quality bases were removed from the sequencing reads using scythe (v0.994) (Buffalo) and sickle (v1.33) (Joshi and Fass), respectively, with default parameters. Filtered sequencing reads were then assembled using SPAdes (v3.7.1) (Bankevich et al., 2012), with default parameters.

### Metagenomic sequencing

#### Skin and Oral

Metagenomic DNA from swabs in lysis buffer and beads was extracted using the GenElute Bacterial DNA Isolation kit (MilliporeSigma) with the following modifications: each sample was digested with 50 μg of lysozyme, and 5 units lysostaphin and mutanolysin for 30 minutes prior to beadbeating in the TissueLyser II (QIAGEN) for 2 x 3 minutes at 30 Hz. Each sample was centrifuged for 1 minute at 15000 x g prior to loading onto the GenElute column. Negative (environmental) controls and positive (mock community) controls were extracted and sequenced with each extraction and library preparation batch to ensure sample integrity.

#### Stool

Approximately 50 microliters of the thawed Omnigene preserved stool sample was added to a microfuge tube containing 350 μL Tissue and Cell lysis buffer (Epicentre) and 100 μg 0.1 mm zirconia beads (BioSpec Products). Metagenomic DNA was extracted using the QiaAmp 96 DNA QiaCube HT kit (Qiagen) with the following modifications: each sample was digested with 50 μg of lysozyme, and 5 units lysostaphin and mutanolysin for 30 minutes prior to beadbeating in the TissueLyser II (QIAGEN) for 2 x 3 minutes at 30 Hz. Each sample was centrifuged for 1 minute at 15000 x g prior to loading 200 ul into an S plate. Negative (environmental) controls and positive (mock community) controls were extracted and sequenced with each extraction and library preparation batch to ensure sample integrity.

Sequencing adapters and low quality bases were removed from the mWGS reads using scythe (v0.994) (Buffalo) and sickle (v1.33) (Joshi and Fass), respectively, with default parameters. Host reads were removed by mapping all sequencing reads to the hg19 human reference genome using Bowtie2 (v2.3.1) (Langmead and Salzberg, 2012; Langmead et al.), under “very-sensitive” mode. Unmapped reads (i.e., microbial reads) were used to estimate the relative abundance profiles of the microbial species in the samples using MetaPhlAn2 (Segata et al., 2012; Truong et al., 2015, analysis date 3/2020).

### Strain typing with Biotyper IR

The Bruker IR Biotyper Fourier Transform Infrared (FT-IR) Spectroscopy system (Bruker Daltonics, Germany) was used to evaluate strain differences between isolates of a given species. The IR Biotyper analyzes the spectra of peaks corresponding to cell surface glycoproteins and uses hierarchical clustering to establish relationships between strains. Strains were grown from single colonies to a state of confluent growth on tryptic soy agar plates (TSA). An overloaded 1 μL inoculating loop of cell material was resuspended in 50 microliters of 70% ethanol (Sigma-Aldrich) in a 1.5 ml Bruker suspension vial with inert metal cylinders, and vortexed to homogeneity. 50 μL of deionized water was added to the tube, and again vortexed. 15 μL of each isolate suspension was pipetted onto 4 spots of a silicon microtiter plate along with 2 spots each of Bruker Infrared Test Standards 1 and 2. The plate was allowed to dry in a 37°C incubator, then loaded into the IR Biotyper for analysis. Spectra were processed by the IR Biotyper software in the 1300-800cm-1 wavelength, corresponding to the carbohydrate region. Each spectra was comprised of 521 different datapoints. For exploratory analysis to assess similarity of spectra, we used the default IR Biotyper software settings to generate principal components analysis (PCA) plots and dendograms via hierarchical clustering using Euclidean distance to generate distance matrices.

### Comparative genomics of microbial genomes

Microbial genomes were prepared using our ‘dirty’ DNA prep, as described previously^17^, where 1 mL of overnight culture was centrifuged at 20,000 x g for 1 minute before the bacterial pellet was resuspended with 100 mL of 1X TE and transferred to a 2 mL bead beating tube with 100-125 mL 0.5 mm diameter glass beads (BioSpec Products). An additional 100 mL of 1X TE was added to the tube, followed by vortexing of the sample for 30 s at max speed (3000 rpm) on a Vortex Adaptor (Mo Bio Laboratories). The mixture was then centrifuged at 13,000 x g for 5 minutes to pellet the cellular debris, and the supernatant was transferred to a new tube to be used as template for Nextera XT library preparation as for metagenomic sequencing.

WGS reads were quality-filtered, trimmed, and assembled as described previously^17^. The resulting draft genomes, as well as publicly available genomes (**Table S1**) were analyzed using the classify workflow (with default parameters) of GTDB-Tk (v1.0.2, reference database version r89)^18^. Based on the bacterial marker gene alignment generated by GTDB-Tk, a phylogenetic tree was inferred using FastTree (v.2.1.11)^19^ with default parameters and visualized using Figtree (v1.4.4)^20^.

### Data deposition

Strains generated herein are available upon reasonable request. Genomes and metagenomic data are deposited in the Short Read Archive (SRA) under Bioproject TBD and Accession TBD.

## RESULTS AND DISCUSSION

Our goal was to identify a core set of cultivation conditions for each body site that would allow relative throughput across individuals to maximize the number of species and strains obtained with reasonable throughput, rather than a comprehensive recovery of microbes from each body site. For the skin, we chose the forehead and toe web space as representatives of an oily and a moist skin site, respectively, and their microbiota differ markedly in our previous surveys^17,21,22^. We chose the inner cheek and tongue dorsum to represent the oral cavity, and stool for gut. Because different individuals can harbor markedly different microbial species and strains^6^, we obtained, collectively, 25 samples from 12 individuals, with at least 5 samples for each body site. For each sample, we then performed shotgun metagenomic sequencing (1.6±1.0×10^6^, 2.0±1.1×10^6^, 10.5±1.0×10^6^ quality-controlled, human DNA dehosted reads for oral, skin, and stool, respectively, **Table S1**) and culturomics as described below.

### Cultivation conditions and species identification

Guided by our previous metagenomic data and a literature search^8,11,12,14-16,23–38^, we compiled the aerobic and anaerobic cultivation conditions reported in **Table 1** and **Figure 1**. To examine the proportion of microbes recovered by these conditions, we first characterized the fungal, bacterial, and viral composition of our samples using shotgun metagenomics as it is culture-independent and yields the most unbiased compositional reconstruction (**Figure 2, Table S2**). Consistent with previous reports, *Cutibacterium acnes* and *Corynebacterium* sp. were most abundant in the oily sites of the forehead, and staphylococci and *Corynebacterium* sp. in the foot. In the cheek, streptococci, *Rothia mucilanginosa*, and *Haemophilus parainfluenzae* were most abundant, and in the tongue dorsum, *Neisseria flavescens, Prevotella* sp., *Veillonella* sp., and streptococci. Finally, in the gut, *Bacteroidales* and *Clostridiales* were most abundant.

**Figure 2.**
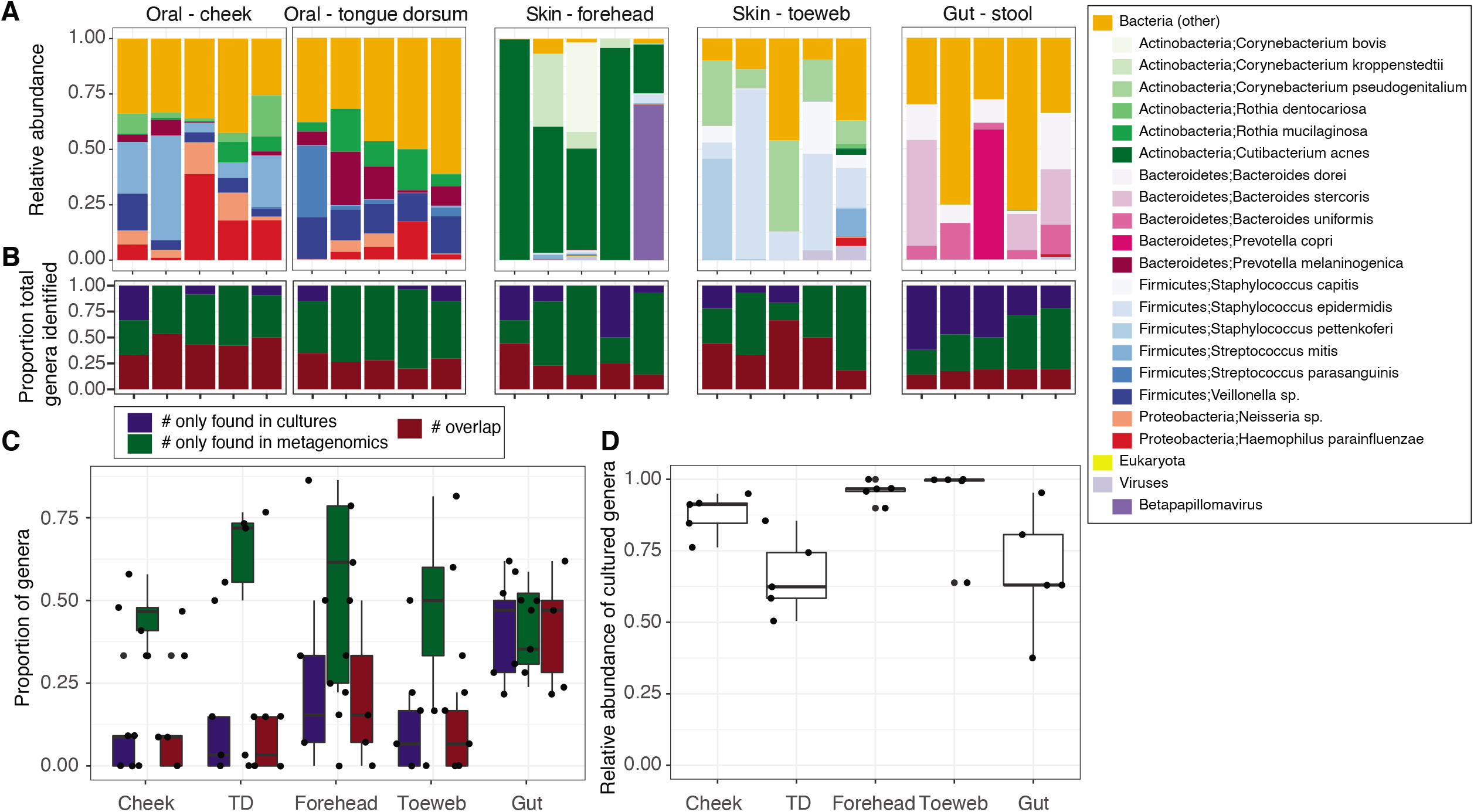
Metagenomic reconstructions of community composition and representation by cultivars. **A)** Relative abundance plots of oral, skin and gut samples from this study; each bar is an individual sample and the top 20 most abundant species are plotted. Lesser abundance bacteria, fungi, and viruses are collectively represented by their respective kingdom. The proportion of bacterial genera cultivated or identified through sequencing for each sample, **B),** or across samples shown by boxplot, **C)**. Red shows proportion of bacterial genera identified through both cultivation and metagenomics, and blue and green show the proportion of bacterial genera identified by only one method. **D)** The total relative abundance of the original metagenomic sample (bacteria only) that is accounted for by the genera cultivated (overlap in **C**).

In culturing, we recommend a rule of thumb to dilute samples 1:10 and 1:100 for skin sites, 1:100 and 1:1000 for oral sites, 1:1000 and 1:10000 for gut samples prior to plating, reflecting the low, medium, and high microbial bioburden of these sites, respectively. Dilution plates usually consisted of 1-4 dominant microbes with singletons interspersed at low density. Because of this we aimed to select ~12 microbes per plate, selecting up to 3 of each visibly unique morphology for subculturing to further purify the selected isolate, followed by MALDI-TOF mass spectrometry analysis for identification. MALDI-TOF accuracy at the species and genus level varies widely by taxonomy and even instrument (e.g., ~84% for species, ~92% for genus in a recent estimate of anaerobic bacteria^39^, but up to 98% accuracy and 94% accuracy can be observed in *Enterobacteriaceae* and staphyloccci^40^), with disease-associated species having the deepest reference databases and thus the highest corresponding accuracy. Overall, we obtained 15 unique genera (34 species) in the skin, 17 genera (53 species) in the mouth, and 41 genera (97 species) from stool from 600, 1155, and 1451 isolates tested, respectively (**Table 2**).

**Table 2.** Genera uniquely detected by culturomics, metagenomics, and overlap

| Phylum | Genus | # samples cultured only | # samples metagenomic only | # overlaps | # total samples |
| --- | --- | --- | --- | --- | --- |
| Actinobacteria | Corynebacterium | 5 | 2 | 12 | 19 |
| Firmicutes | Streptococcus | 1 | 7 | 11 | 19 |
| Firmicutes | Veillonella | 0 | 7 | 10 | 17 |
| Proteobacteria | Neisseria | 0 | 4 | 10 | 14 |
| Actinobacteria | Actinomyces | 2 | 3 | 10 | 15 |
| Actinobacteria | Rothia | 0 | 6 | 9 | 15 |
| Firmicutes | Staphylococcus | 8 | 0 | 9 | 17 |
| Actinobacteria | Cutibacterium | 1 | 3 | 8 | 12 |
| Proteobacteria | Haemophilus | 0 | 9 | 7 | 16 |
| Firmicutes | Gemella | 0 | 7 | 6 | 13 |
| Firmicutes | Lachnoanaerobaculum | 0 | 5 | 5 | 10 |
| Bacteroidetes | Bacteroides | 0 | 1 | 5 | 6 |
| Bacteroidetes | Parabacteroides | 0 | 0 | 5 | 5 |
| Fusobacteria | Fusobacterium | 1 | 3 | 4 | 8 |
| Firmicutes | Lactobacillus | 2 | 2 | 3 | 7 |
| Actinobacteria | Bifidobacterium | 0 | 1 | 3 | 4 |
| Actinobacteria | Collinsella | 0 | 0 | 3 | 3 |
| Bacteroidetes | Prevotella | 0 | 12 | 2 | 14 |
| Proteobacteria | Aggregatibacter | 0 | 6 | 2 | 8 |
| Firmicutes | Clostridium | 3 | 4 | 2 | 9 |
| Actinobacteria | Micrococcus | 2 | 2 | 2 | 6 |
| Proteobacteria | Escherichia | 3 | 1 | 2 | 6 |
| Actinobacteria | Brevibacterium | 1 | 1 | 2 | 4 |
| Firmicutes | Flavonifractor | 2 | 0 | 2 | 4 |
| Firmicutes | Granulicatella | 0 | 13 | 1 | 14 |
| Actinobacteria | Atopobium | 1 | 5 | 1 | 7 |
| Actinobacteria | Kocuria | 1 | 5 | 1 | 7 |
| Firmicutes | Eubacterium | 1 | 3 | 1 | 5 |
| Proteobacteria | Acinetobacter | 0 | 3 | 1 | 4 |
| Bacteroidetes | Alistipes | 0 | 3 | 1 | 4 |
| Firmicutes | Blautia | 1 | 2 | 1 | 4 |
| Bacteroidetes | Odoribacter | 1 | 2 | 1 | 4 |
| Actinobacteria | Kytococcus | 1 | 1 | 1 | 3 |
| Firmicutes | Bacillus | 5 | 0 | 1 | 6 |
| Actinobacteria | Dermabacter | 2 | 0 | 1 | 3 |
| Bacteroidetes | Capnocytophaga | 0 | 9 | 0 | 9 |
| Proteobacteria | Actinobacillus | 0 | 8 | 0 | 8 |
| Fusobacteria | Leptotrichia | 0 | 7 | 0 | 7 |
| Bacteroidetes | Porphyromonas | 0 | 7 | 0 | 7 |
| Bacteroidetes | Alloprevotella | 0 | 7 | 0 | 7 |
| Firmicutes | Oribacterium | 0 | 6 | 0 | 6 |
| Firmicutes | Stomatobaculum | 0 | 6 | 0 | 6 |
| Firmicutes | Subdoligranulum | 0 | 6 | 0 | 6 |
| Proteobacteria | Campylobacter | 2 | 5 | 0 | 7 |
| Firmicutes | Peptostreptococcus | 2 | 5 | 0 | 7 |
| Firmicutes | Megasphaera | 0 | 5 | 0 | 5 |
| Firmicutes | Solobacterium | 0 | 5 | 0 | 5 |
| Firmicutes | Oscillibacter | 0 | 5 | 0 | 5 |
| Proteobacteria | Enhydrobacter | 0 | 4 | 0 | 4 |
| Firmicutes | Abiotrophia | 0 | 4 | 0 | 4 |
| Proteobacteria | Kingella | 0 | 4 | 0 | 4 |
| Firmicutes | Dorea | 0 | 4 | 0 | 4 |
| Proteobacteria | Bilophila | 0 | 4 | 0 | 4 |
| Firmicutes | Faecalibacterium | 0 | 4 | 0 | 4 |
| Firmicutes | Finegoldia | 2 | 3 | 0 | 5 |
| Firmicutes | Selenomonas | 0 | 3 | 0 | 3 |
| Saccharibacteria | Saccharibacteria | 0 | 3 | 0 | 3 |
| Proteobacteria | Parasutterella | 0 | 3 | 0 | 3 |
| Firmicutes | Roseburia | 0 | 3 | 0 | 3 |
| Firmicutes | Anaerococcus | 3 | 2 | 0 | 5 |
| Firmicutes | Ruminococcus | 2 | 2 | 0 | 4 |
| Firmicutes | Parvimonas | 1 | 2 | 0 | 3 |
| Actinobacteria | Eggerthella | 3 | 1 | 0 | 4 |
| Firmicutes | Pediococcus | 2 | 1 | 0 | 3 |
| Firmicutes | Enterococcus | 6 | 0 | 0 | 6 |
| Proteobacteria | Citrobacter | 3 | 0 | 0 | 3 |
| Actinobacteria | Dietzia | 3 | 0 | 0 | 3 |
| Firmicutes | Lysinibacillus | 3 | 0 | 0 | 3 |

We then sought to examine the degree to which these isolates represented the predicted composition from metagenomic data. At the genus level (to account for MALDI-TOF and metagenomic classification accuracy at the species level), we observed an overlap of cultivated and metagenomic genera of 44.4±7.7% (cheek, mean ± standard deviation), 27.9±5.4% (tongue dorsum), 24.1±12.4% (forehead), 24.1±18.0% (toeweb), and 18.1±2.3% (stool, **Figure 2B-C**). These represented a high proportion of the community composition reconstructed with metagenomic sequencing (**Figure 2D**); 87.7±7.5%, 66.2±13.8%, 95.8±3.7%, 92.6±16.1%, and 67.9±21.7%, respectively, suggesting that our methods captured the majority of the abundant genera irrespective of body site and that the missing genera were likely low abundance microbes.

We then examined if, and what genera were preferentially cultivated and found there were genera that were identified uniquely by metagenomic sequencing but also uniquely by cultivation (**Table 2, Figure 2B-C**). Metagenomics, as expected, definitively recovered a much greater number of total microbes that were not captured by culturomics (**Figure 2B**). Prevalent metagenomic genera that were never cultivated were primarily anaerobes from the gut and oral cavity, including Bacteroidetes *Capnocytophaga* (found in 9 samples), *Porphyromonas* (7), *Alloprevotella* (7), Proteobacteria *Actinobacillus* (7), and Fusobacteria *Leptotrichia* (7). We were also surprised that we consistently identified species that were not captured by metagenomics. Interestingly, Firmicutes *Enterococcus* was never identified in metagenomic data but fairly extensively cultivated as several species *(avium, faecalis, faecium* across six samples). Similarly, *Bacillus* species (*circulans, pumilus, subtilis*) were only identified in one metagenomic sample (as *Bacillus amyloliquefaciens*) compared to five samples for culturomics. Even well-studied bacteria like staphylococci could be cultivated from a sample more frequently than detected by metagenomic data; 8/17 times it was only detected by cultivation. Potential explanations for these observations could include: 1) species are insufficiently abundant to be classified by the use of clade-specific marker genes as we have performed^41^ but are easily cultivatable (e.g., staphylococci, *Enterococcus)* or 2) incomplete or few reference genomes are available for that genus to enable its classification (e.g., *Bacillus*, but not the case for *Enterococcus* which had hundreds of reference genomes).

Finally, we note two general observations. First, we observed significantly different recovery rate for many microbes. For example, staphylococci in the skin, *E. coli* and *Enterococcus* in the gut, and streptococci in the oral cavity are recovered with far greater frequency and repetition than more abundant but more fastidious microbes. Second, we found that most microbes could be recovered on multiple growth conditions, and that there were relatively few media that specifically allow recovery of a desired taxon. For example, selective staphylococcal media (e.g., SaSelect, Biorad) is designed for colorimetric differentiation of staphylococci, but frequently recovers *Bacillus* and *Micrococcus*. Anecdotally, we have explored depletion of such microbes like staphylococci by antibiotics, lysostaphin, and crystal violet, which result in low-to-moderate depletion, and per the goals of the project at hand, are recommended for further exploration. Overall, we believe that our approach results in broad recovery of abundant bacteria and low-abundance, non-fastidious bacteria. We recommend on a per-application basis, evaluation of targeted approaches for the recovery of desired microbes, either via depletion of abundant microbes, increased number of growth conditions, and potentially most importantly, increased numbers of colonies surveilled. In addition, some microbes are epibionts, requiring co-culture for growth^e.g.,10^. In addition, an increasing number of innovative approaches, including engineered antibody capture^10^, microfluidics devices that can be placed in the environment^9^, 3-D organoids to better recapitulate growth environments, and high throughput content screenings, are being developed to facilitate increased recovery of desired microbes for follow-up experimentation.

### Strain identification

An emerging frontier of metagenomic discovery is the understanding of strain biology, as microbial diversity is ultimately manifested at this finest taxonomic resolution where individual strains of a microbial species can exhibit widely diverse phenotypes. As our methods cultivate dominant microbes, it is particularly effective for investigating strain variation between individuals and cohorts, as we can reliably isolate overlapping species from a given sample (e.g., staphylococci, *E. coli*). Further, extensive strain variation can exist not only between, but as we have shown, within individuals. This phenomena has major implications for disease severity^17^. It is thus valuable to be able to rapidly differentiate strains to understand strain diversity, to identify disease-causing strains, and to prioritize strains for phenotyping.

Different methods with widely differing resolution have been developed for strain typing primarily for clinical use, perhaps most commonly multi-locus sequence typing (MLST), which sequences polymorphisms in highly conserved genes to bin strains into sequence ‘types’. The gold standard is whole genome sequencing, but despite extraordinary technical advances, it remains relatively costly and slow to perform and analyze on large scales. Fourier-transform infrared spectroscopy (FT-IR) can rapidly generate discriminatory biochemical fingerprints primarily based on cell surface macromolecules, e.g., lipids, proteins, and carbohydrates. It has been used in examining clonal outbreaks of varying origin, although overall it remains less frequently used despite its lengthy technical history (reviewed in ^42^), likely because of incomplete understanding of the link between genetic diversity and cell surface macromolecular diversity.

Here, we evaluated the ability of the Bruker IR Biotyper to rapidly differentiate genetically diverse strains from phylogenetically diverse species, selected as common species of interest in the skin, oral, or gut microbiota. In addition to cultivars obtained in our study, we included additional publicly available, fully sequenced isolates to provide additional genetic diversity. Finally, by way of benchmark, we sequenced, or obtained from public repositories, the genomes of these strains to determine genetic relatedness (**Table S1**), although we recognize that the cell surface macromolecules are encoded and modified by numerous genetic pathways and environmental conditions, like length of growth time and composition of growth media, and are likely difficult to translate to genetic distance, as previously noted^42^.

We investigated IR’s ability to differentiate genetically diverse strains of *S. aureus, S. epidermidis, C. acnes* as important skin microbes, and *E. coli* and *B. subtilis* from the gut. We primarily investigated diverse isolates obtained from different individuals, with the exception of *E. coli* in which we investigated within-individual diversity (or clonality) by typing multiple isolates obtained from 4 individuals, each with at least 3 technical replicates (i.e., multiple ‘spots’ of the same colony). To identify general concordance between the phylogenetic distance (genomics, **Figure 3A**) and biochemical distance (IR) between isolates for each species, we performed an exploratory analysis comparing dendrograms (**Figure 3C**) and principal components analysis (**Figure 3B**) generated from both datatypes.

**Figure 3.**
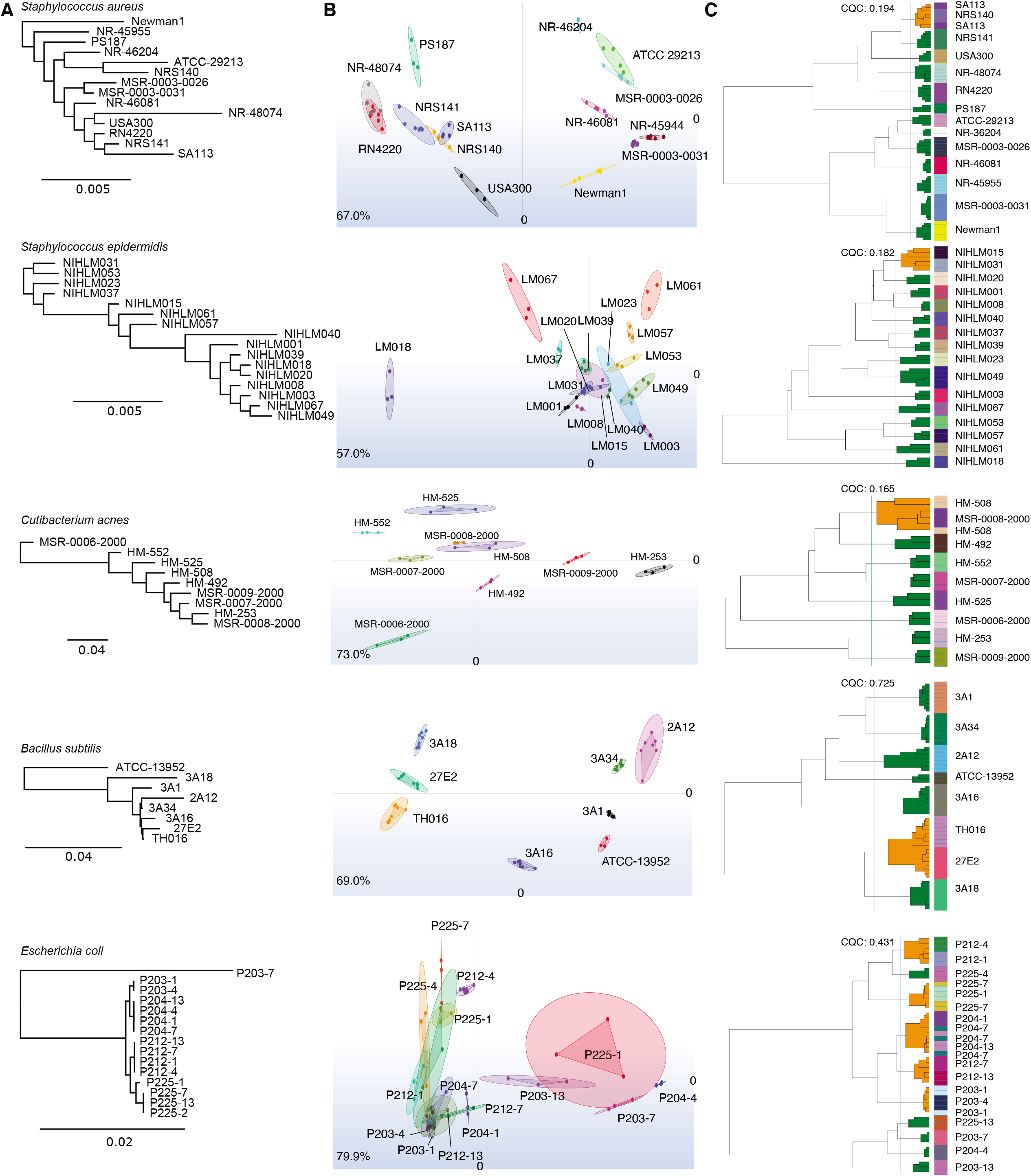
Biotyper IR differentiates genetically distinct strains. **A)** Phylogenetic trees of strain genomes tested in the Biotyper IR analysis based on alignment of bacterial marker genes. Genetic distance is shown in dendrogram; genomes used (generated in this study or obtained from public repositories) are in **Table S1**. **B)** Principal components analysis (PCA) plot showing clustering of strains for each species, with each color representing a unique isolate and each dot within that color representing the isolate’s replicate spectral measurements. Links to the dots showing the variance of the technical replicates; output from IR Biotyper interface. ##% in lower left corner indicate the sum of variance explained by the first two principal components. **C)** Dendrogram of isolates based on spectral measurements; output from IR Biotyper interface. Green and orange in dendrogram represent cluster purity as determined by the Bruker IR software, based on technical replicates of strain spectra: green (“GOOD”), orange (“BAD”). Cluster quality criterion (CQC) indicates how well replicate measurements of an isolate cluster with themselves as well as the purity or homogeneity of each cluster.

Nearly universally, technical replicates were the most similar to each other, irrespective of species and supporting IR’s reproducibility. For each species, we give some examples of concordance and discordance of clusters formed by IR vs. genome sequencing (**Figure 3**).

1. In most cases, *S. aureus* strains formed distinct clusters; in particular, *S. aureus* PS187, Newman1, and NR-45944 which were also identified as more distant phylogenetically. However, in cases where overlapping clusters were identified by IR (e.g., MSR-0003-0026 & ATCC 29213, SA113 & NRS140, and RN4220 & NR-48074), most were relatively genetically distinct. In some cases, genetic and IR clusters were recapitulated (NRS141 & SA113, NR-46204 & ATCC-29213). Surprisingly, MSR-0003-0031 and MSR-0003-0026 were both highly genetically similar (isolated from the same individual) but did not form overlapping clusters.
2. We observed two clades of *S. epidermidis* strains, consistent with our previous large-scale genome analysis^17^. However, these clades were not recapitulated by IR, with the most distinct IR clusters coming from more genetically related strains (NIHLM018, NIHLM067, NIHLM061). Even highly genetically similar strains (e.g., NIHLM018 and NIHLM020), formed distant and distinct clusters.
3. *C. acnes* strains, each isolated from different individuals, largely formed distinct clusters with the exception of MSR-0006-2000 and HM-508, shared overlapping clusters, but were distant genetically. HM-253 and MSR-0008-2000 were genetically most similar but were more closely related to other strains by IR spectra.
4. The *B. subtilis* strains tested had relatively higher genetic distance than for the previous species. TH016 and 27E2, which were most similar genetically, formed overlapping IR clusters, but 3A18, which was relatively more distant, was also a near neighbor. ATCC-13952 robustly formed a distinct cluster and was an outlier genetically. In the case of *B. subtilis*, we also tested biological replicates (independent colonies between two separate runs), which yielded highly concordant results with strains from both runs grouping into the same cluster(s).
5. The ability to differentiate *E. coli*, where we examined both within- and between-individual variation, differed from the other species tested in its relatively lower clarity in strain differentiation by IR. However, upon closer look, in most cases, strains from the same patient clustered together based on their spectral profile (e.g., P212-4 & P212-1, P225-4 & P225-1 & P225-7, P204-1 & P204-7 & P204-13, P203-1 & P203-4). Because *E. coli* within an individual likely derives from a single lineage (like *B. fragilis* in the gut^43^), this underscores a strength of the IR in its ability to discriminate clonal strains from other strains.

A surprising result was 203-7, which had been identified as *E. coli* by MALDI-TOF, and which genetically was relatively distant from the other *E. coli* strains (taxonomic identification based on alignment of a set of core marker genes), including other strains from the same patient, but shared an IR spectral cluster with another patient’s strain (P225-13). Upon closer examination, this strain, which had been identified as *E. coli* by MALDI-TOF, was classified as *Klebsiella pneumoniae*, which shares 99.01% ANI in the single copy marker genes used for classification. It is possible that species-level variation may, at times, exceed strain variation or even variation between technical replicates. Anecdotally, we have observed similar results when testing different strains from different species on the IR (data not shown). This reinforces the need for strain isolation coupled with a multipronged approach to characterization.

Collectively, these results suggest that IR’s ability to differentiate strains is first, likely species-specific, as different species can have notably different levels of strain-level genetic diversity (i.e., some bacteria have relatively closed vs. open pangenomes, e.g., *C. acnes*^44^ and *S. epidermidis*^17^, respectively) that may differentially affects cell surface macromolecule diversity. Second, IR may be more valuable in differentiating clonal strains from any other non-clonal strain, rather than making a general assessment of genetic divergence and phylogenetic placement. In addition, it is important to note additional limitations of the IR. For example, run-to-run reproducibility in differentiating the same set of strains is acceptable (data not shown), but it cannot directly compare, for example, two different sets of strains from two different runs as analysis is within-run, rather than benchmarked to an external database. Thus, given the 96-spot format, ~30 strains can be typed simultaneously accounting for technical triplicates. Second, as we previously noted, real genetic distance is difficult to deconvolute without, we believe, an extensive paired comparative genomics approach. Nevertheless, despite these limitations, we believe that this rapid strain differentiation would be useful for selecting a subset of isolates from a set of patient cultivars that minimizes likely phenotypic redundancy. For example, *Bacteroides fragilis* has been deemed relatively clonal in the gut^43^, and thus selecting just one representative isolate per individual might suffice, but for *S. epidermidis*, which has extensive within-individual strain diversity^17^, several representative strains might be chosen.

In conclusion, here we have presented and characterized a facile workflow for cultivation of bacteria from skin, oral, and gut microbiota from genus/species to the strain level with a focus on a high level of throughput across many patient samples rather than comprehensive recovery of all the species within a sample. We believe these efforts add to an increasing technical collection of approaches for metagenomic discovery, and highlights the value-add of strain-level analysis to better understand genetic and phenotypic diversity underlying host-microbiome interactions.

---

**Table S1. Metagenomic read count statistics and genomes used in the study**

**Table S2. Relative abundances of bacteria, fungi, and viruses in metagenomic data**

## Consent for publication

All authors have consented to publication.

## Author contributions

Conceptualization: JO. Methodology: EF, VP, AH. Analysis: EF, VP, AH, WZ, RH, YON, JD, RB, AP, JO. Writing: JO, EF, AH. Resources: JO.

## Acknowledgements

We thank the Oh lab for critical feedback and the Microbial Genomics Services and Genome Technologies Cores at the Jackson Laboratory for technical assistance. We also thank Michael Santino from Bruker for critical reading of the manuscript.

